# OneD: increasing reproducibility of Hi-C Samples with abnormal karyotypes

**DOI:** 10.1101/148254

**Authors:** Enrique Vidal, François le Dily, Javier Quilez, Ralph Stadhouders, Yasmina Cuartero, Thomas Graf, Marc A. Martí-Renom, Miguel Beato, Guillaume J. Filion

**Affiliations:** Gene Regulation, Stem Cells and Cancer Program, Centre for Genomic Regulation (CRG), The Barcelona Institute of Science and Technology (BIST), Dr. Aiguader 88, 08003, Barcelona, Spain; Universitat Pompeu Fabra (UPF), Barcelona, Spain; CNAG-CRG, Centre for Genomic Regulation (CRG), Barcelona Institute of Science and Technology (BIST), Baldiri i Reixac 4, 08028 Barcelona, Spain; ICREA, Pg. Llus Companys 23, 08010 Barcelona, Spain

## Abstract

The three-dimensional conformation of genomes is an essential component of their biological activity. The advent of the Hi-C technology enabled an unprecedented progress in our understanding of genome structures. However, Hi-C is subject to systematic biases that can compromise downstream analyses. Several strategies have been proposed to remove those biases, but the issue of abnormal karyotypes received little attention. Many experiments are performed in cancer cell lines, which typically harbor large-scale copy number variations that create visible defects on the raw Hi-C maps. The consequences of these widespread artifacts on the normalized maps are mostly unexplored. We observed that current normalization methods are not robust to the presence of large-scale copy number variations, potentially obscuring biological differences and enhancing batch effects. To address this issue, we developed an alternative approach designed to take into account chromosomal abnormalities. The method, called *OneD*, increases reproducibility among replicates of Hi-C samples with abnormal karyotype, outperforming previous methods significantly. On normal karyotypes, *OneD* fared equally well as state-of-the-art methods, making it a safe choice for Hi-C normalization. *OneD* is fast and scales well in terms of computing resources for resolutions up to 1 kbp. *OneD* is implemented as an R package available at <u>http://www.github.com/qenvio/dryhic.</u>

## 1 Introduction

One of the crown achievements of modern biology was to realize that genomes have an underlying three-dimensional structure contributing to their activity (Rowley and Corces, 2016; Dekker and Mirny, 2016; Pezic *et al.*, 2017). In mammals, this organization plays a key role in guiding enhancer-promoter contacts (De Laat and Duboule, 2013), in V(D)J recombination (Choi and Feeney, 2014) and in X chromosome inactivation (Galupa and Heard, 2015). A significant breakthrough towards this insight was the development of the high throughput chromosomal conformation capture technology (Hi-C), assaying chromosomal contacts at a genome-wide scale (Lieberman-Aiden *et al.*, 2009). Nowadays, exploring the spatial organization of chromatin has become a priority in many fields and Hi-C has become part of the standard molecular biology toolbox (Dekker *et al.*, 2013).

Contrary to the precursor technologies 3C, 4C and 5C (Dekker *et al.*, 2002; Simonis *et al.*, 2006; Dostie *et al.*, 2006; de Wit and de Laat, 2012), Hi-C interrogates all possible pairwise interactions between restriction fragments. However, this does not guarantee that the method has no bias. On the contrary, local genome features such as the G+C content, the availability of restriction enzyme sites and the mappability of the sequencing reads have been shown to impact the results (Yaffe and Tanay, 2011), in addition to general experimental biases such as batch effects. It is thus important to normalize Hi-C data in order to remove biases and artifacts, so that they are not confused with biological signal.

Several methods have been proposed to remove biases in Hi-C experiments (Schmitt *et al.*, 2016). The first strategy is to model biases explicitly from a defined set of local genomic features, such as the G+C content. This approach is used in the method of Yaffe and Tanay (2011) and in Hicnorm by Hu *et al.* (2012). The second strategy is to implicitly correct unknown biases by enforcing some regularity condition on the data. This approach is used in the Iterative Correction and Eigenvector decomposition method *(ICE)* of Imakaev *et al.* (2012), whereby the total amount of contacts of every bin is imposed to be the same. *ICE* is currently the most popular method, due in part to its speed.

Neither of these strategies were designed for cell types with karyotypic aberrations, most common in cancer. Yet, Hi-C is very sensitive to aneuploidy, copy number variations and translocations. Actually, these aberrations have so much influence on the outcome that they can be used as signatureto re-assemble the target genome (Korbel and Lee, 2013). An additional complication is that karyotypic aberrations are not experimental biases, so it is unclear whether they should be corrected at all or be considered part of the biological signal.

So far, the only attempt to address the issue was the chromosome-adjusted Iterative Correction Bias method *(caICB)* of Wu and Michor (2016). However, *caICB* applies a uniform chromosome-wide copy number correction, effectively excluding the numerous cases of partial aneuploidy and regional copy number variations.

Here we propose *OneD*, a method to correct local chromosomal abnormalities in Hi-C experiments. *OneD* explicitly models the contribution of known biases via a generalized additive model. The normalized data is more reproducible between replicates and across different protocols. Importantly, *OneD* is also efficient when cells have a normal karyotype, where it performs as well as the best normalization methods. Finally, the implementation is as fast as *ICE* and it scales up to 1 kbp resolution with reasonable computing resources.

## 2 Methods

### 2.1 Model

The most common representation of Hi-C data is a contact matrix, obtained by slicing the genome in *n* consecutive bins of fixed size (the resolution) and computing the number of contacts between each pair of bins. The values are stored in the cells of the contact matrix (*x_ij_*), quantifying the interaction between the two loci at positions *i* and *j.*

Our approach is to model the tally of contacts for each bin, thus reducing the matrix to a one dimension score (hence the name *OneD).* We assume that the total number of contacts per bin (*t_i_*) can be approximated by a negative binomial distribution. This choice is sensible because the amount of contacts is a discrete variable and because the negative binomial distribution allows for overdispersion. We further assume that the explicit sources of bias have independent contributions to the mean of the distribution for a given bin (*λ_i_*).

Given that this relationship might not be linear (see for instance Figure 1A), we allowed a smooth representation using thin plate penalized regression splines (Wood, 2003) in a generalized additive model (Wood, 2011). The model can be parametrized as

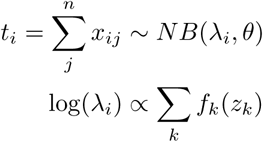

**Figure 1:**
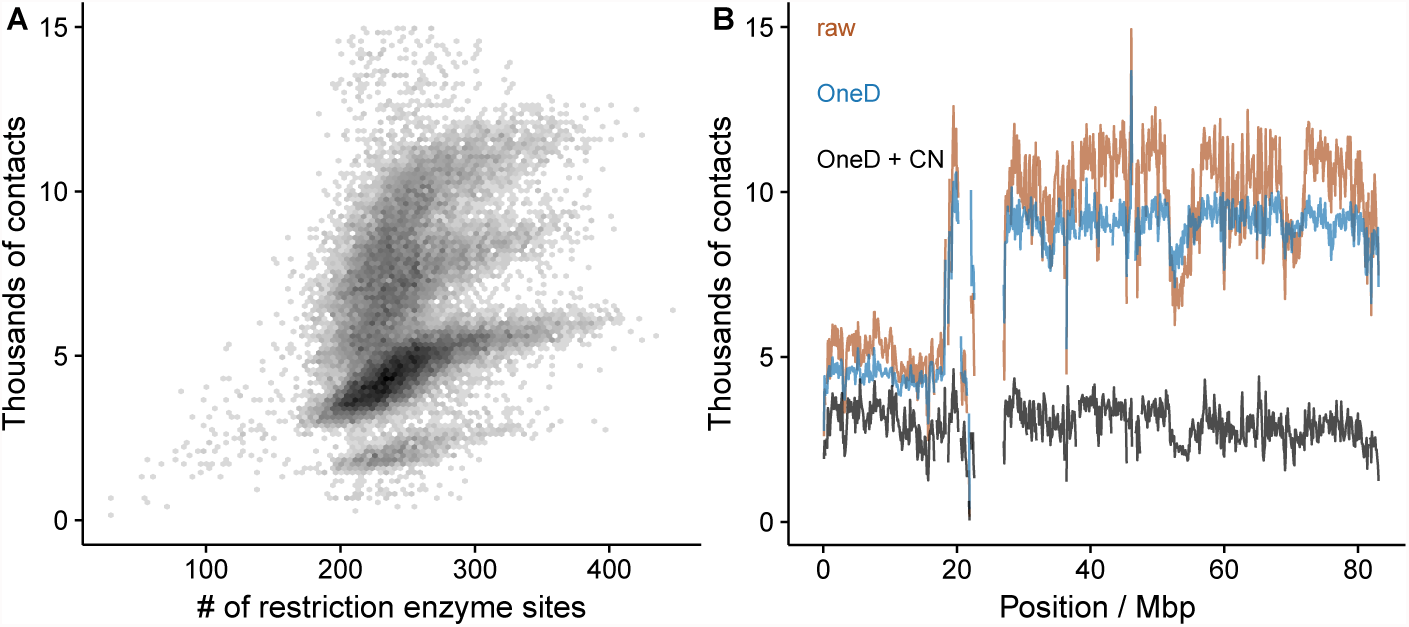
Total amount of contacts per bin. A. Non linear relationship between the number of restriction sites and the total number of contacts per bin in T47D. B. Total number of contacts per bin on chromosome 17 of T47D. The brown line represents the raw signal, the blue line represents the signal after bias correction, the black line represents the signal after bias and copy number (CN) correction. The long arm of chromosome 17 (the region corresponding to 20-80 Mbp) is present in four copies, explaining that the signal is about twice higher than for the short arm.

Where *x_ij_* is the raw number of contacts between bins *i* and *j*, and *z_k_* is the additive bias of genomic feature *k.* The smooth functions {*f_k_*(·)} are estimated jointly with the negative binomial dispersion parameter *θ* using the `mgcv` package (Wood, 2011) of the R software (R Core Team, 2017).

Once the parameters of the model are determined, the estimated means {λ_i_} are rescaled to obtain a correction vector 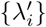 that can be used to compute the corrected counts 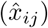.

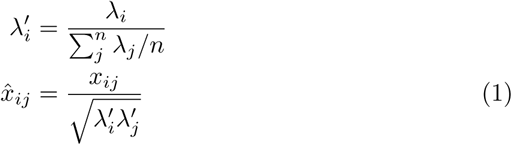

In line with previous methods (Yaffe and Tanay, 2011; Hu *et al.*, 2012), the default features used to fit the model are the local G+C content, the read mappability and the content and number of restriction sites. The model and the implementation can be modifed or extended with any user-provided genomic features.

### 2.2 Copy number correction

Briefly, a hidden Markov model with emissions distributed as a Student’s *t* variable is fitted on the corrected total amount of contacts per bin 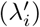. The model consists of 7 states that correspond to 1, 2, 3, 4, 5, 6 and 8 copies of the target, for a total of 3 emission parameters (a single scaling parameter, a single standard deviation and a single degree of freedom for all the states) and 21 transition parameters.

The model is fitted with the Baum-Welch algorithm (Baum and Petrie, 1966) until convergence, following a previously described implementation (Filion *et al.*, 2010). The Viterbi path is then computed and corresponds to the inferred copy number of each bin (*c_i_*).

A correction equal to the square root of the copy number is then applied to the whole matrix. More specifically, each cell is updated to

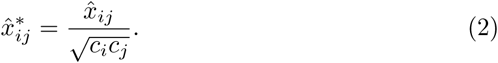

### 2.3 Data sources

To test the correction of biases, we gathered a set of published (Le Dily *et al.*, 2014; ENCODE Project Consortium, 2012; Rao *et al.*, 2014; Stadhouders *et al.*, 2017; Lin *et al.*, 2012; Dixon *et al.*, 2012) and unpublished Hi-C data of different cell types and organisms (Table 2.3).

We used several experiments comprising T47D breast cancer cell lines (7 samples), K562 leukemia cell lines (4 samples), both with aberrant karyotypes; and mouse primary B cells (6 samples) and ES cells (7 samples), both with normal diploid karyotypes. The experiments were carried out in different laboratories, following either the original Hi-C protocol (Lieberman-Aiden *et al.*, 2009) or the newer *in situ* version (Rao *et al.*, 2014), and using different restriction enzymes (DpnII, HindIII, MboI and NcoI). We also used array-based copy-number segmentation of the two cell lines obtained from the COSMIC database (Forbes *et al.*, 2010).

**Table 1:** Hi-C data sets used in this study. Sample ID: Unique sample identifier in the respective projects. Cell type: source of the biological sample. Application: Hi-C protocol. RE: Restriction enzyme used for the digestion. Sequencing core: Laboratory performing the experiment (CRG: Centre for Genomic Regulation; UMass: University of Massachusetts; Baylor: Baylor College of Medicine; UCSD: University of California San Diego; LICR: Ludwig Institue for Cancer Research). Set(s): Sets in which the sample was included (see text for detail). Source: SRA, GEO or ENCODE raw data identifier.

| Sample ID | Cell type | Application | RE | Sequencing core | Set(s) | Source |
| --- | --- | --- | --- | --- | --- | --- |
| dc3a1e069_51720e9cf | T47D | *in situ* Hi-C | DpnII | CRG | T47D, K562 | NA |
| b1913e6c1_51720e9cf | T47D | *in situ* Hi-C | DpnII | CRG | T47D | NA |
| dc3a1e069_ec92aa0bb | T47D | *in situ* Hi-C | DpnII | CRG | T47D, K562 | NA |
| HindIII-T0 | T47D | dilution Hi-C | HindIII | CRG | T47D | SRR1054341 |
| NcoI-T0 | T47D | dilution Hi-C | NcoI | CRG | T47D | SRR1054343 |
| ENCLB758KFU | T47D | dilution Hi-C | HindIII | UMass | T47D | ENCLB758KFU |
| ENCLB183QHG | T47D | dilution Hi-C | HindIII | UMass | T47D | ENCLB183QHG |
| HIC069 | K562 | *in situ* Hi-C | MboI | Baylor | T47D, K562 | SRR1658693 |
| HIC070 | K562 | *in situ* Hi-C | MboI | Baylor | T47D, K562 | SRR1658694 |
| HIC071 | K562 | *in situ* Hi-C | MboI | Baylor | K562 | SRR1658695,SRR1658696 |
| HIC074 | K562 | *in situ* Hi-C | MboI | Baylor | K562 | SRR1658701,SRR1658702 |
| b7fa2d8db_bfac48760 | B cell | *in situ* Hi-C | DpnII | CRG | mm10 | GSE96611 |
| fc3e8b36a_7bf1bf374 | ES cell | *in situ* Hi-C | DpnII | CRG | mm10 | GSE96611 |
| b7fa2d8db_7284b867a | B cell | *in situ* Hi-C | DpnII | CRG | mm10 | GSE96611 |
| fc3e8b36a_38bfd1b33 | ES cell | *in situ* Hi-C | DpnII | CRG | mm10 | GSE96611 |
| b7fa2d8db_58e812fc2 | B cell | *in situ* Hi-C | DpnII | CRG | mm10 | GSE96611 |
| fc3e8b36a_c990a254e | ES cell | *in situ* Hi-C | DpnII | CRG | mm10 | GSE96611 |
| b7fa2d8db_73f11d923 | B cell | *in situ* Hi-C | DpnII | CRG | mm10 | GSE96611 |
| fc3e8b36a_4bf044f18 | ES cell | *in situ* Hi-C | DpnII | CRG | mm10 | GSE96611 |
| GSM987817 | B cell | dilution Hi-C | HindIII | UCSD | mm10 | SRR543428-SRR543431 |
| GSM987818 | B cell | dilution Hi-C | HindIII | UCSD | mm10 | SRR543432-SRR543442 |
| GSM862720 | ES cell | dilution Hi-C | HindIII | LICR | mm10 | SRR443883-SRR443885 |
| GSM862721 | ES cell | dilution Hi-C | HindIII | LICR | mm10 | SRR400251-SRR400256 |
| GSM862722 | ES cell | dilution Hi-C | NcoI | LICR | mm10 | SRR443886-SRR443888 |

All data were processed through a pipeline based on TADbit (Serra *et al.*, 2016). Briefly, after controlling the quality of FASTQ files, paired-end reads were mapped to the corresponding reference genome (hg38 and mm10) taking into account the restriction enzyme site. Non-informative contacts were removed applying the following TADbit filters: `self-circle, dangling-ends, error, extra-dangling-ends, duplicated and random-breaks.` For more details, see the methods section of Stadhouders *et al.* (2017). In addition, the pipeline is available from the supplementary material published by Quilez *et al.* (2017).

We developed the routines contained in the `dryhic` R package to efficiently create sparse representations of contact matrices and further apply *vanilla*, *ICE* and *oneD* corrections. `HiTC` (Servant *et al.*, 2012) and HiCapp (Wu and Michor, 2016) were used to carry out the *LGF* and *caICB* corrections respectively. All the results are based on a resolution of 100 kbp, but we found no major differences at different resolutions (not shown).

### 2.4 Comparison of Hi-C matrices

There is no universally accepted standard to compare Hi-C matrices. The simplest metric is the Spearman correlation applied to intra-chromosomal contacts up to a given distance (5 Mb in what follows). The second option is to measure the similarity of observed over expected contacts via the Pearson correlation up to a given distance range. Compared to the first, this metric gives more weight to changes occurring away from the diagonal. The third option is to compute a correlation per distance stratum and then obtain a stratum-adjusted correlation coefficient (SCC) as defined by Yang *et al.* (2017). Finally, the last option, proposed by Yan *et al.* (2017) is to measure the Pearson correlation between the last eigenvectors of the Laplacian of the Hi-C matrix. This approach borrows the concepts of spectral clustering (Von Luxburg, 2007) and amounts to comparing high level features of the matrix.

We defined three data sets to measure experimental reproducibility after normalization: The first contained the samples from T47D plus two samples from K562, the second contained the samples from K562 plus two samples from T47D, the third contained all the mouse samples (see Table 2.3 for details). Given a set of experiments and a metric, we first computed all pairwise combinations between experiments and then classified the comparisons according to the characteristics of each pair (cell type, protocol, batch and treatment).

To measure the gain or loss of similarity upon normalization, we compared raw matrices to obtain a baseline. The differences with this baseline were estimated using a linear mixed model fitted with the `lmer` function of the `lme4` R package (Bates *et al.*, 2015), where the fixed effect was the normalization method and the random effect was the chromosome. Receiver operating characteristic (ROC) curves were computed using the `ROCR` package (Sing *et al.*, 2005).

## 3 Results

### 3.1 Experimental bias correction

The principle of *OneD* is to explicitly model Hi-C biases on a single variable: the total amount of contacts for each bin of the matrix (see 2.1 for detail). The reason for this choice is that the total amount of contacts is approximately proportional to the local copy number. For instance, a duplicated region in a diploid genome will show on average a 50% increase in the number of contacts. Discontinuities of the amount of contacts thus correspond to changes of the copy number.

Experimental biases affect the total amount of contacts in a continuous but not necessarily linear way. Figure 1A shows the relationship between the amount of contacts and the number of restriction enzyme sites in T47D, a breast cancer cell line with an aberrant karyotype. Four clouds are visible. Each corresponds to sequences present in one to four copies. In all of them, the relationship flattens as the number of restriction sites increases. To capture this relationship, *OneD* fits a non-linear model between the total amount of contacts and the known sources of bias (by default the G+C content, the number of restriction sites and the mappability of the reads).

The experimental biases are estimated genome-wide and each cell of the matrix is then corrected using equation (1). Note that the corrected amount of contacts is still proportional to the copy number. Figure 1B shows the corrected number of contacts along chromosome 17 of T47D. *OneD* greatly reduces the wiggling of the total amount of contacts (blue line).

In what follows, we benchmarked *OneD*, against *vanilla*, *ICE* (Imakaev *et al.*, 2012), *caICB* (Wu and Michor, 2016) and the Local Genomic Features method *(LGF*, Hu *et al.*, 2012; Servant *et al.*, 2012). The first three methods correct biases implicitly, whereas the fourth method does it explicitly.

Given that our model is based on total number of contacts, we reasoned that a preliminary test would be to check if the corrected number of contacts per bin reflects the copy number (as measured by an independent technique such as array-based copy-number segmentation). We tested the validity of this approach against the Catalogue Of Somatic Mutations In Cancer (COSMIC, Forbes *et al.*, 2010). Figure 2 shows the Pearson correlation between the corrected number of contacts and the copy number estimation for T47D and K562 (a leukemia cell line with an aberrant karyotype). Similar results were obtained using Spearman correlation (Supplementary Figure 1). All the methods except *OneD* decreased the agreement between the signal and the copy number because they partially corrected the discrepancy. In contrast, *OneD* enhanced the conformity of the signal with the copy number. Not correcting for variable copy number at that stage may seem counter-intuitive, but the tests below will show this leads to better performance.

**Figure 2:**
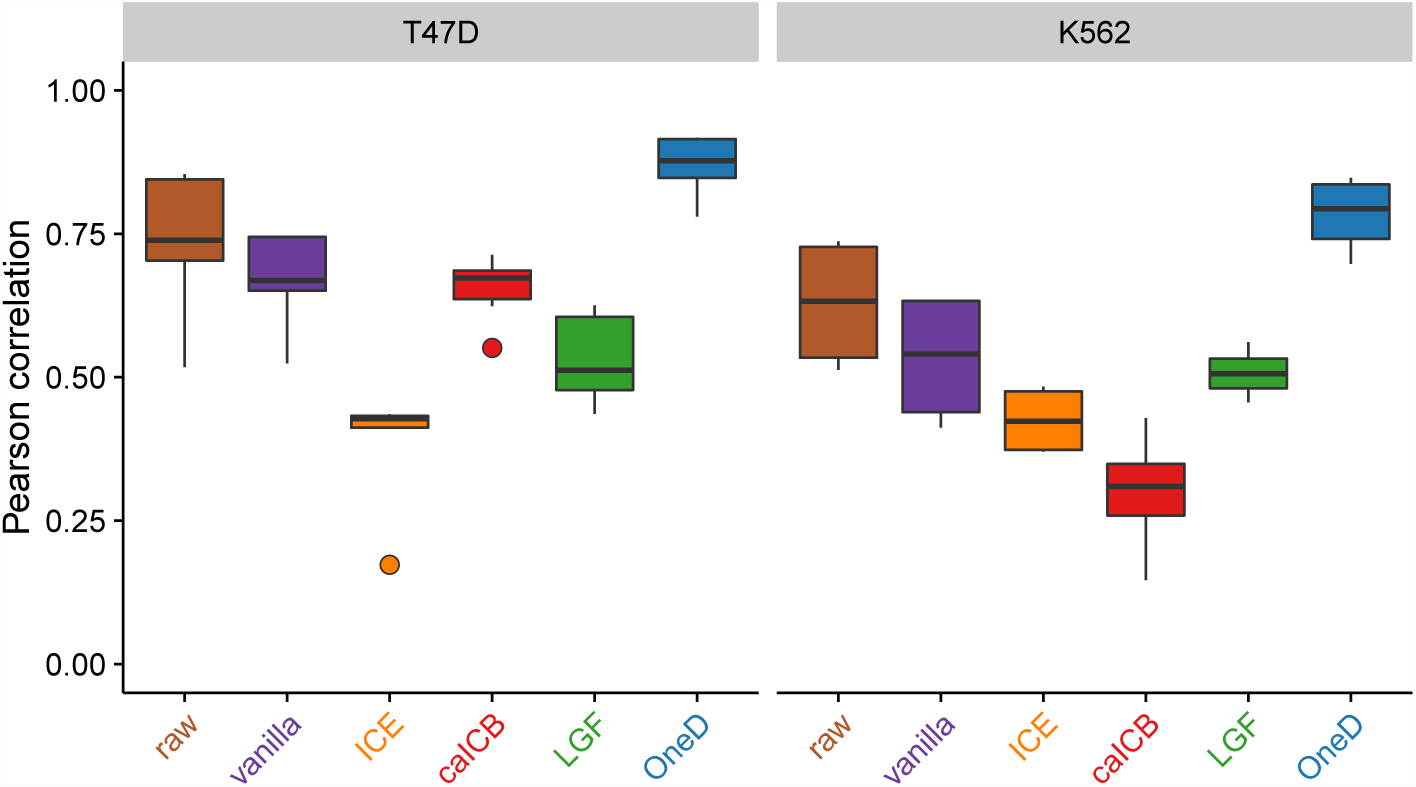
Pearson correlation between the total number of contacts per bin and an independent estimation of the copy number (COSMIC). Left panel T47D breast cancer cell line, right panel K562 leukemia cell line.

### 3.2 Aberrant karyotypes

We first benchmarked the performance of our approach on biological samples with an aberrant karyotype. A good normalization method should increase the similarity between biological replicates by reducing irrelevant experimental variance, such as batch effects, laboratory of origin and protocol variations. Similarly, a good normalization should decrease the similarity between different samples to enhance the biological differences.

We assembled two Hi-C data sets obtained from T47D and K562 cells. In each set we spiked two samples from the other cell line (Table 2.3) to introduce biological variability. We compared matrices before and after normalization by different methods using the Pearson correlation of observed over expected counts (see 2.4). This gave an indication of the impact of a given normalization method. The results are summarized in Figure 3.

**Figure 3:**
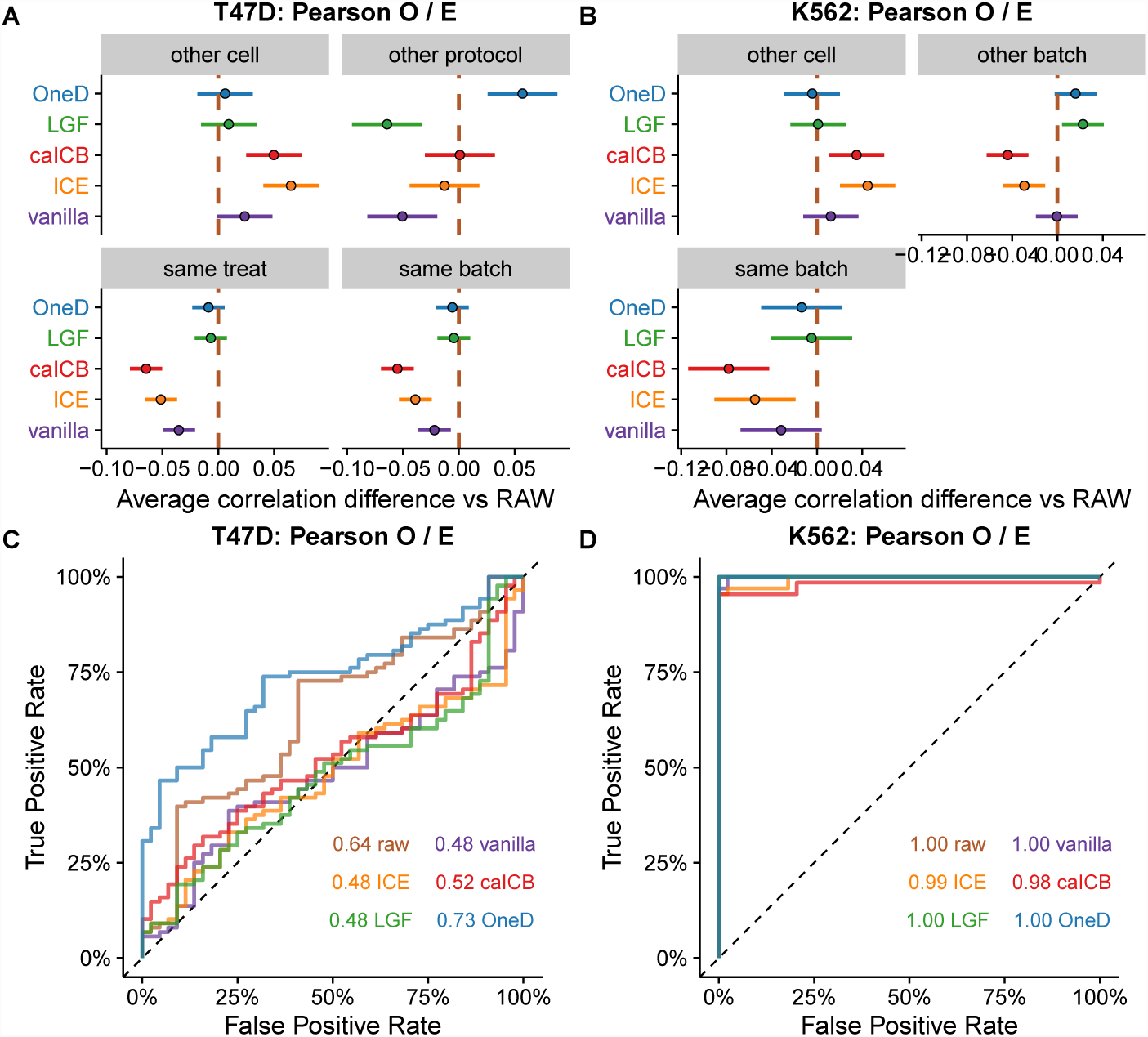
Results of the comparison between samples with aberrant karyotypes. A and B. Average changes compared to the raw data on the T47D and K562 sets. The bars represent 95% confidence intervals centered on the mean difference of the correlation score between a given correction method and the raw data. C and D. ROC curves on the T47D and K562 sets. The areas under the curve are indicated in the bottom right corner. The color code is the same as panels A and B. The brown lines correspond to raw matrices. All results in this figure are based on Pearson correlations between the observed over expected counts.

The *caICB* and *ICE* methods increased the similarity between the different cell lines (Figures 3A and 3B and Supplementary Figures 2 and 3). This is an undesirable effect, as it obscures the biological variability. Likewise, these methods decreased the similarity between samples that received the same treatment (Figure 3A), suggesting that the normalization process is detrimental to the biological signal in these two cases. The method *vanilla* followed the same trend but to a lesser extent, consistent with the fact that it consists of a single *ICE* iteration. *OneD* was the only method to increase the similarity between experiments carried out on the same material but with a different protocol (Figure 3A).

An important application of normalization methods in experimental setups is to identify outliers. We thus investigated the capacity of the different methods to help identify the samples from the other cell type spiked in the data set. We interpreted the pairwise comparison scores as classifier scores and summarized the results with a ROC curve (Figures 3C and D). All the methods, including the absence of normalization, succeeded in identifying the T47D outliers among the K562 samples, but recognizing the K562 outliers among the T47D samples proved more challenging. *OneD* increased the discrimination power compared to the raw matrices, but all the other methods decreased it. Actually, they performed little better than random on this task. Using the other metrics described in 2.4 yielded similar results (Supplementary Figures 5, 6 and 7). Note that Spearman correlation of contacts presented the worst performance for all scenarios, and it thus seems to be a poor choice for a metric to compare Hi-C matrices.

Taken together, these results show that correcting experimental biases with *OneD* enhances the biological variation and reduces the noise on samples with an aberrant karyotype.

### 3.3 Normal karyotypes

Does the performance of *OneD* on aberrant karyotypes come at the cost of decreased performance on normal karyotypes? To address this question, we assembled another data set comprised of mouse B cells and embryonic stem (ES) cells, both with a normal karyotype. The cell types were pooled in almost equal proportions (see Table 2.3) and the same tests as above were performed.

In these conditions, the experimental protocol had a strong effect on the impact of the different normalization methods (Figure 4A). For instance, *caICB* and *ICE* increased the similarity when the protocols were different, but decreased it when the protocols were the same. The effect was stronger when comparing identical cell types, but the same trend appeared when comparing different cell types, indicating that these methods may enhance or reduce biological variation, depending on the context. Once more, *vanilla* followed the same trend as *ICE* but to a lesser extent. The *LGF* method increased the similarity when comparing the same cells with different experimental protocols, and had little to no effect in the other cases. This indicates that *LGF* is a safe choice in this case.

**Figure 4:**
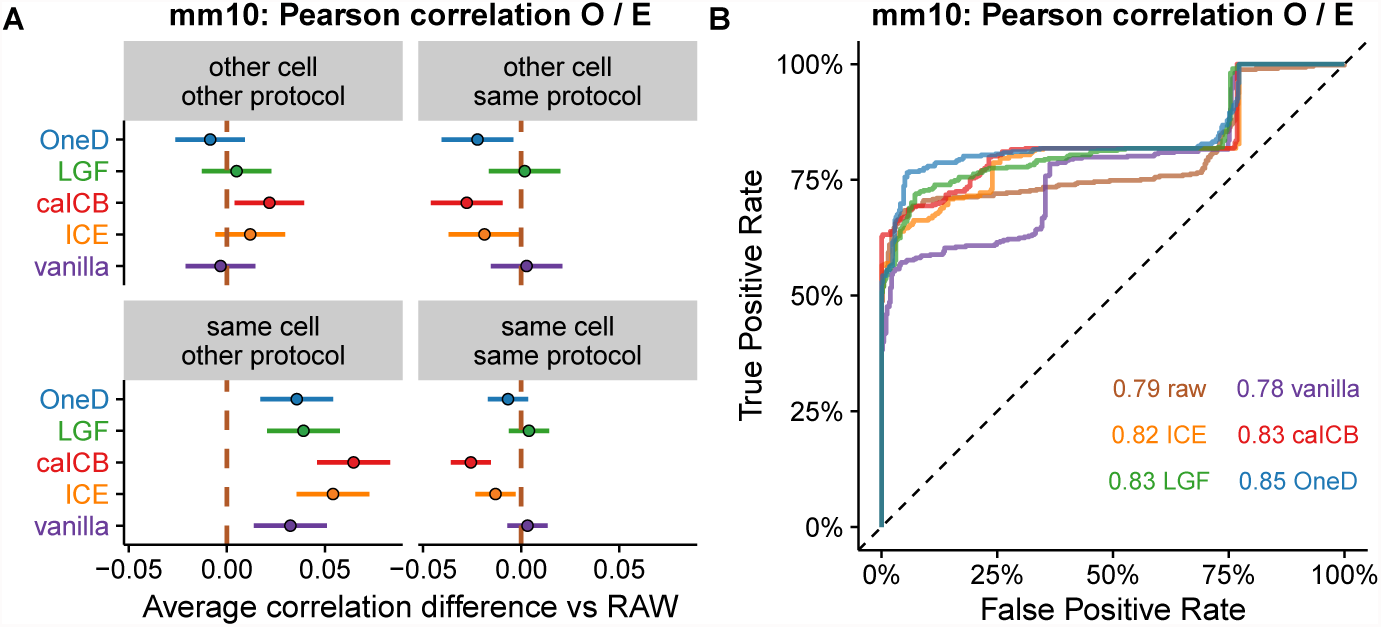
Results of the comparison between samples with normal karyotypes. A. Average changes compared to the raw data on the mouse data set. B. ROC curves on the mouse data set. The legend is as in Figure 3.

*OneD* decreased the similarity between different cell types when using the same protocol and increased it between identical cell types when using different protocols. In the other two cases, it had little effect. In summary, *OneD* never enhanced the experimental noise and even reduced it in one more case than *LGF.*

When interpreting the similarity scores as classification scores, we observed that all the methods could identify approximately 50% of the biological pairs, after which their performance diverged (Figure 4B). *OneD* achieved the highest area under the curve on this problem, but with a small margin over the other methods except *vanilla.* Using other metrics to compare matrices gave similar results: *OneD* was always among the top scoring methods (Supplementary Figures 7 and 8). In these conditions, Spearman correlation of contacts again appeared as the worst comparison metric because it showed a lower performance for all the normalization methods.

Taken together, these results indicate that *OneD* performs as well as the best normalization methods, even with normal karyotypes.

### 3.4 Copy number correction

Removing the explicit experimental biases enhances the importance of the copy number in the signal (Figure 2). The copy number is not an experimental bias, but it may be considered an additional source of bias to be removed. For this reason, *OneD* also allows the user to correct the copy number and output an euploid-equivalent matrix. To do so, *OneD* segments the linear signal of the total amount of contacts into piecewise homogeneous regions. This is carried out by a hidden Markov model whereby the averages of the states are constrained to be an integer number, up to a scaling factor (see 2.2). This allows the model to detect regions with a number of copies equal to 1, 2, 3 *etc.* With the inferred number of copies at hand, *OneD* then normalizes each cell of the matrix with equation (2).

Different normalization methods can either enhance or diminish the signal in regions with higher copy number (Figure 5). In this example, *ICE* over-compensated the original bias at the center of the picture and faded the signal almost entirely. Concomitantly, the signal at the bottom left of the matrix was enhanced and showed a structure that was not visible in the raw data. On the contrary, *LGF* strengthened the central region and the diagonal. *OneD* reduced the level of the central portion by a factor 2 approximately, but did not otherwise distort the main features of the region. This example shows that the copy number does not have a simple and predictable effect on the final matrix. Not taking it into account may open the door to some normalization artifacts.

**Figure 5:**
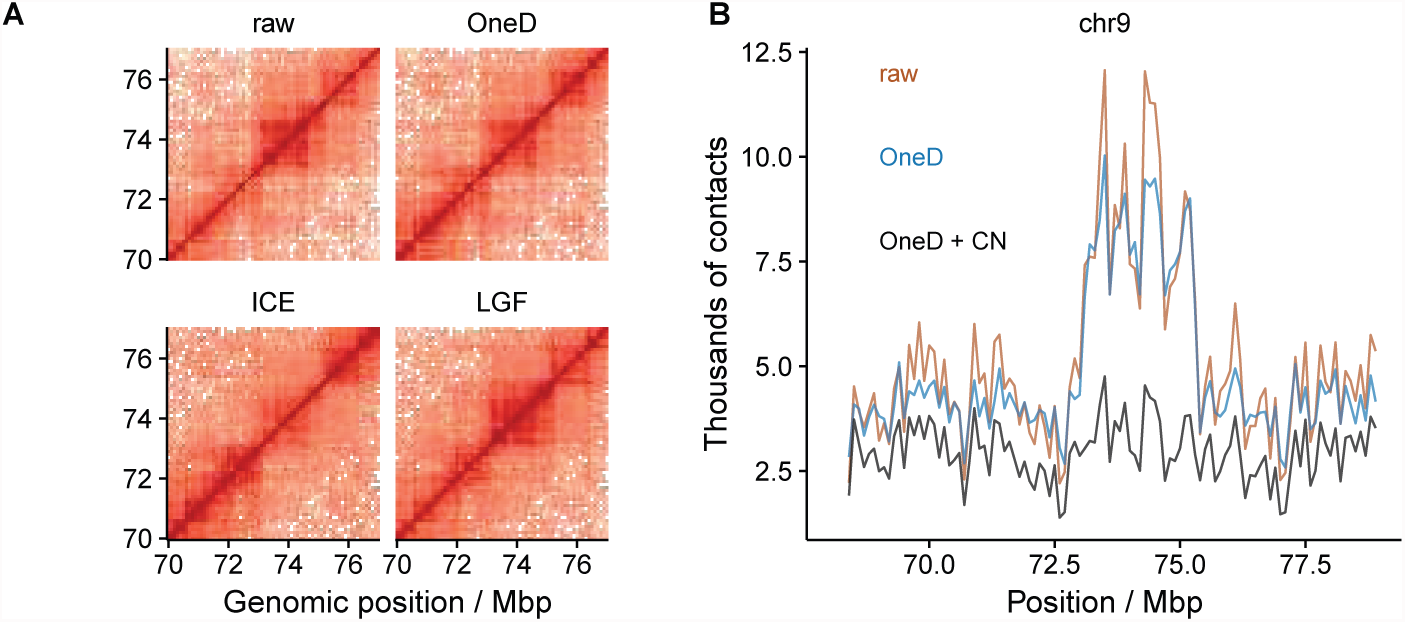
Copy number correction A. Detail of a Hi-C matrix normalized with different methods. The central portion has an increased copy number, which affects normalization. *ICE* fades it away, *LGF* enhances it and *OneD* reduces the signal by about half. B. Profile of the total amount of contacts after copy numer correction. The plot shows the same region as panel A. The brown line represents the raw signal, the blue line represents the signal after bias correction, the black line represents the signal after bias and copy number (CN) correction.

Figure 5B shows the total sum of contacts after correction for the copy number with *OneD.* The region around 72.5-75.0 Mbp showed an elevated amount of contacts in the raw data. After copy number correction, the signal is brought to the same level as the flanking regions. The result provided by *OneD* is not necessarily the right one (see Discussion), but at least it does not correct copy number variations as a side effect of some other criteria.

In summary, *OneD* can be used to obtain an euploid-equivalent normalized matrix in cases where the effect of the copy number must be removed from the signal.

### 3.5 Speed

Finally, we compared the computational speed of the different normalization methods. *vanilla* and *ICE* have broad acceptance for their conceptual simplicity, ease of use and speed (Imakaev *et al.*, 2012). This is even more significant as current explicit methods (Servant *et al.*, 2012) are much slower in comparison.

Unlike the other methods, *OneD* corrects a single variable instead of the whole matrix, and thus estimates the model parameters much faster than previous explicit methods. We measured the running time of the different tools on a 3.3 GHz machine with 62 GB RAM, always using the default parameters. Figure 6A shows the running times of the different methods on the samples described in Table 2.3 at 100 kbp resolution. The fastest method was *vanilla* and the slowest is *LGF*, with an over 100-fold span between the two.

**Figure 6:**
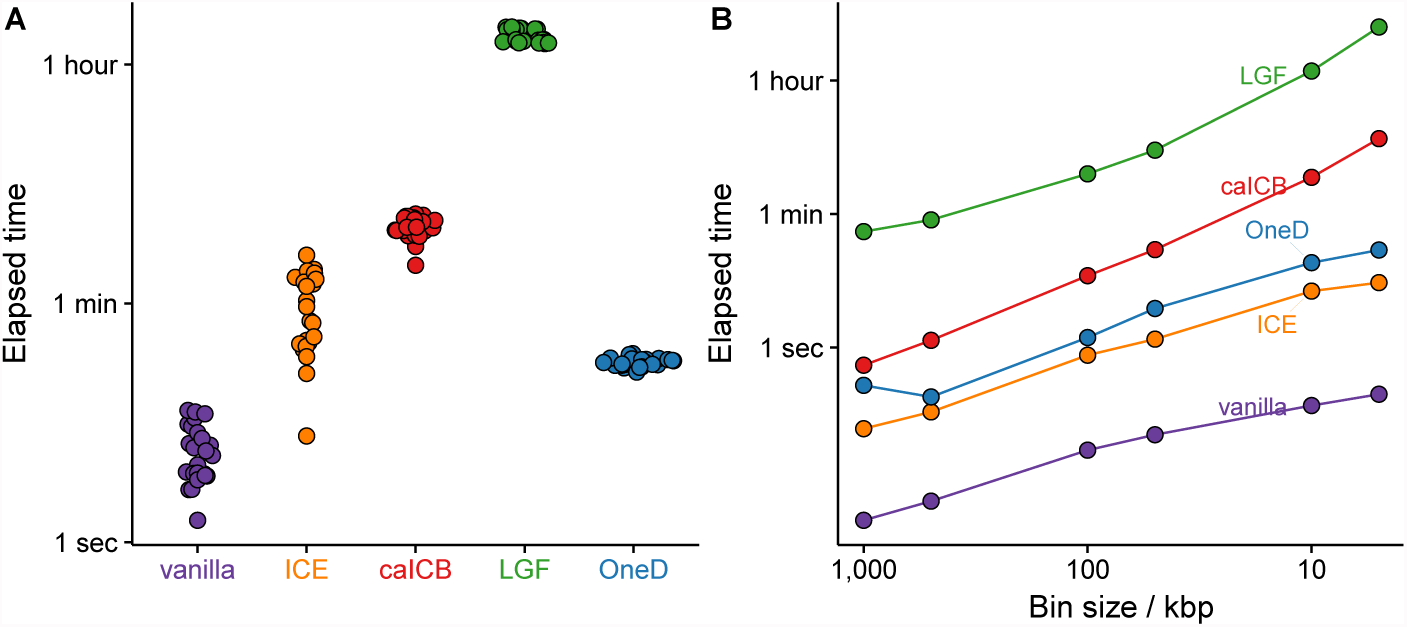
Computing time of the bias correction methods. A. Total time for the entire genome at 100 kbp resolution. Each dot corresponds to one sample. The only method faster than ours under performs in all sample comparisons. B. Time to correct a reduced genome (chr19-22) of one sample at different resolutions. Note the logarithmic scale on the y-axis on both panels.

*OneD* was the second fastest method and it always ran in less than 1 min in the conditions of the benchmark. Throughout this benchmark, the memory footprint of *OneD* was less than 300 MB. Interestingly, the running time of *OneD* was much more homogeneous than that of the other methods. The reason is that the size of the regression problem to be solved by the mgcv package is always the same at fixed resolution.

We also performed a benchmark on a smaller data set at increasing resolution (Figure 6B). On this benchmark, *ICE* ran slightly faster than *OneD*, and the the rank of the methods remained the same at all resolutions. Taken together, these results show that *OneD* is competitive in terms of computational speed compared to existing methods.

## 4 Discussion and Conclusions

Here we introduced *OneD*, a fast computational method to normalize Hi-C matrices. *OneD* was developed ground up to address the need to normalize data from biological samples with aberrant karyotypes, but it applies seamlessly to the case of normal karyotypes. We showed that *OneD* performs significantly better than other methods when the cells present karyotypic aberrations (Figure 3), and that it performs equally well as the best methods on euploid genomes (Figure 4). We also showed that *OneD* is approximately as fast as *ICE*, which makes it competitive from the point of view of computational speed.

The originality of *OneD* lies in that it projects all the biases onto a single variable: the total amount of contacts per bin. This allows greater running speed, while preserving a good performance on samples with karyotypic aberrations. One of the reasons why *OneD* is able to better highlight the biological distinctions between samples is that it only corrects the copy number if specifically requested by the user. The impact of copy number variations on normalization is rather opaque, especially if they are treated as implicit biases (Figure 5). Treating them as explicit biases with optional removal seems to be an overall safer strategy.

This raises the question whether variations of the copy number constitute a biological signal or an artifact. If the biological sample contains karytoypic aberrations, then its genome is grossly different from the reference genome, which makes signal correction very challenging. The proper approach would be to use the actual genome of the biological sample as a reference to construct the contact map. However, this strategy is presently unfeasible because assembling mammalian genomes is still a hard problem.

Depending on the intention of the user, the effect of the copy number should either be kept or removed. This is why *OneD* does not perform the correction by default, but allows the user to obtain a euploid-equivalent Hi-C map computed from a hidden Markov model. The resulting matrices have a near constant amount of contacts per bin, but the artifacts caused by the mismatch between the genome of the sample and the reference genome are still present (for instance, the artifacts caused by large scale inversions are not changed in any way).

Overall, *OneD* constitutes a novel computational approach to normalize Hi-C matrices. If the karyotype of the sample is aberrant, it enhances the biological variation. If not, it performs at least equally well as other methods in terms of quality and of computational speed.

## Acknowledgements

We would like to thank the members of the 4DGenome Synergy project for the fruitful discussions during project meetings. E.V. wants to acknowledge the members of Miguel Beato’s laboratory for their insights during lab meetings.

## Author contributions

Conceptualization, E.V.; Methodology, E.V. and G.F.; Software, E.V. and G.F.; Formal Analysis, E.V.; Investigation, E.V., F.D., Y.C. and R.S.; Resources, M.B.; Data curation, J.Q., Writing Original Draft, E.V. and G.F., Writing Review & Editing, F.D. J.Q., R.S., Y.C, T.G., M.A.M-R. and M.B.; Visualization, E.V.; Supervision, G.F. and M.B., Project Administration, M.B.; Funding Acquisition, M.B., T.G., M.A.M-R. and G.F.

---

**Figure S1:**
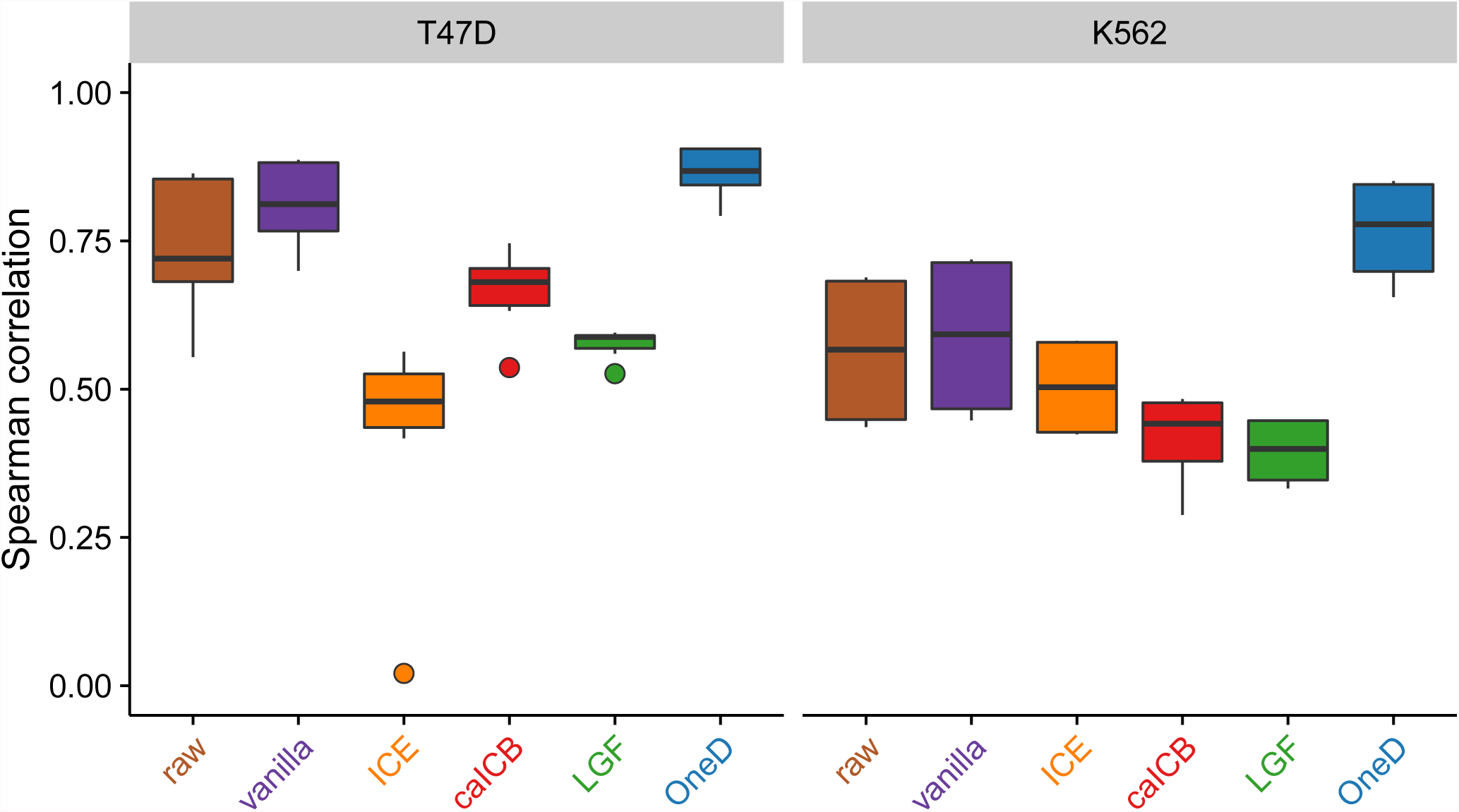
Spearman correlation between total number of contacts per bin and independent copy number estimation (COSMIC) for each of the methods compared. Left panel T47D breast cancer cell line, right panel K562 leukemia cell line. The new proposal (in blue) outperforms the rest of alternatives.

**Figure S2:**
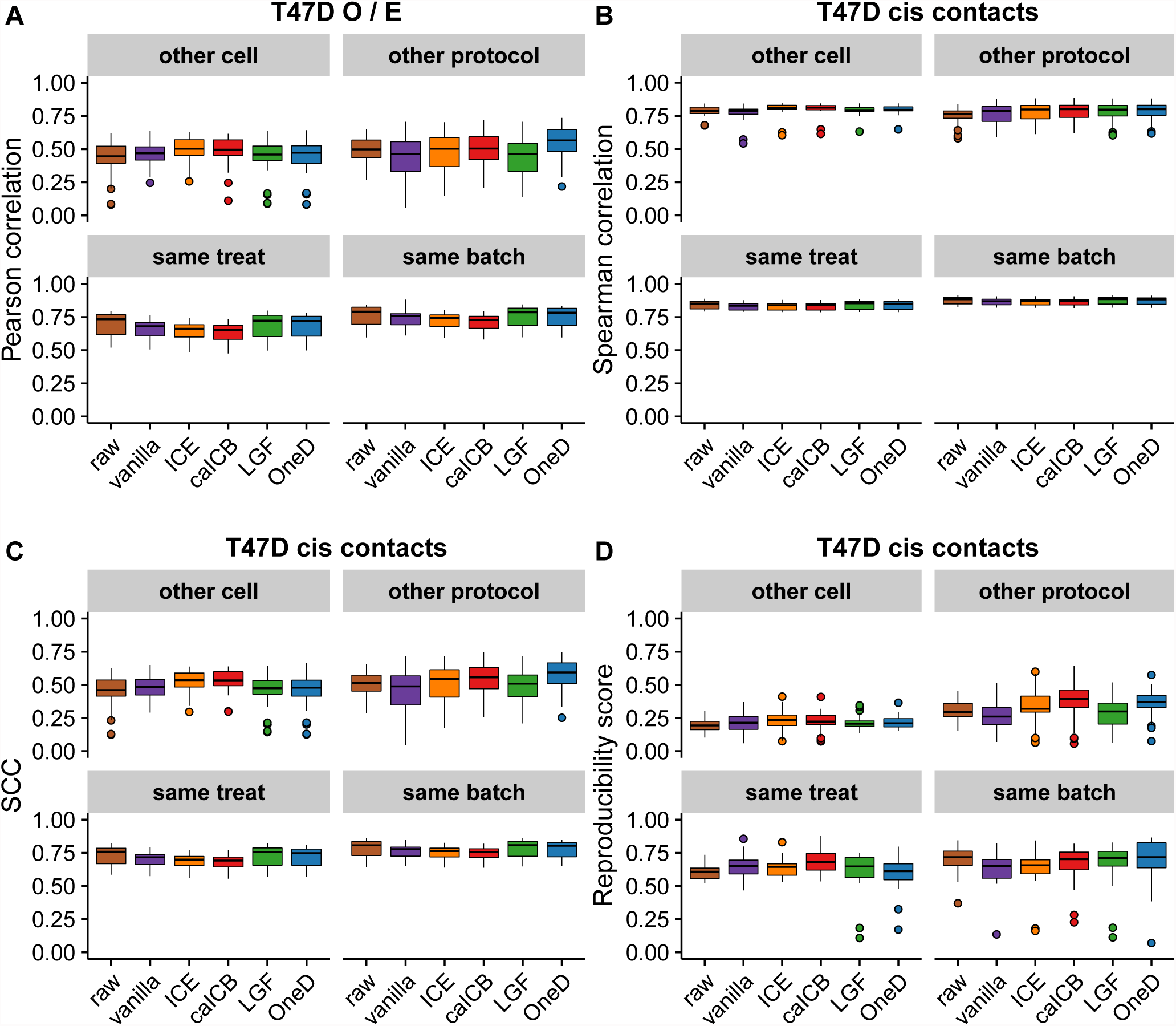
Boxplots representing the results of the pair-wise comparisons of the samples included in the T47D set. X axis: different correction methods. Y axis: correlation. Each panel groups paris of samples with the correspondind characteristics (in terms of cell type, protocol, batch and treatment). A. Pearson correlation between observed over expected counts. B. Spearman correlation between observed counts. C. Stratum-adjusted correlation coefficient (SCC) between observed counts. D. Reproducibility score of observed counts.

**Figure S3:**
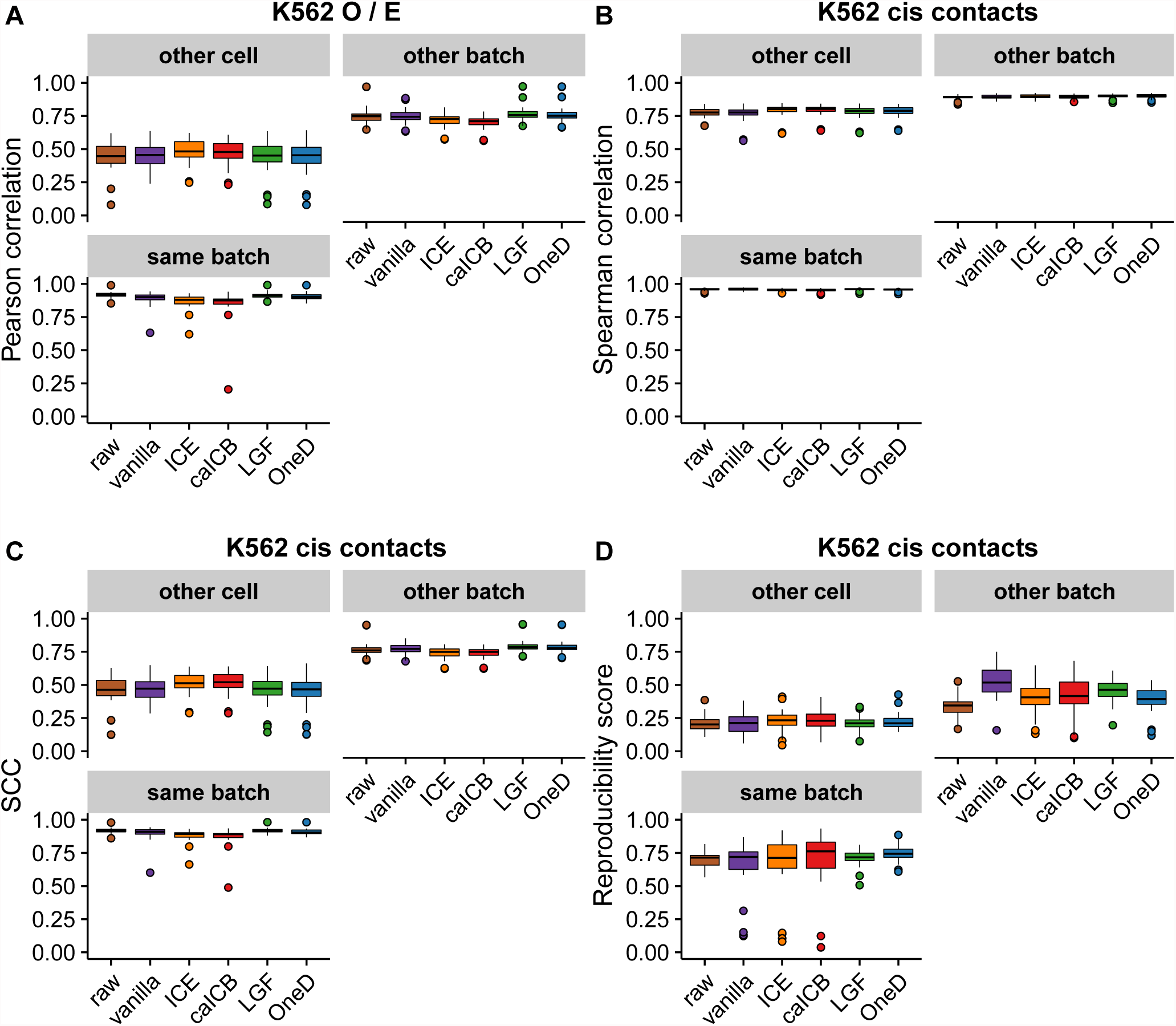
Boxplots representing the results of the pair-wise comparisons of the samples included in the K562 set. X axis: different correction methods. Y axis: correlation. Each panel groups paris of samples with the correspondind characteristics (in terms of cell type, protocol, batch and treatment). A. Pearson correlation between observed over expected counts. B. Spearman correlation between observed counts. C. Stratum-adjusted correlation coefficient (SCC) between observed counts. D. Reproducibility score of observed counts.

**Figure S4:**
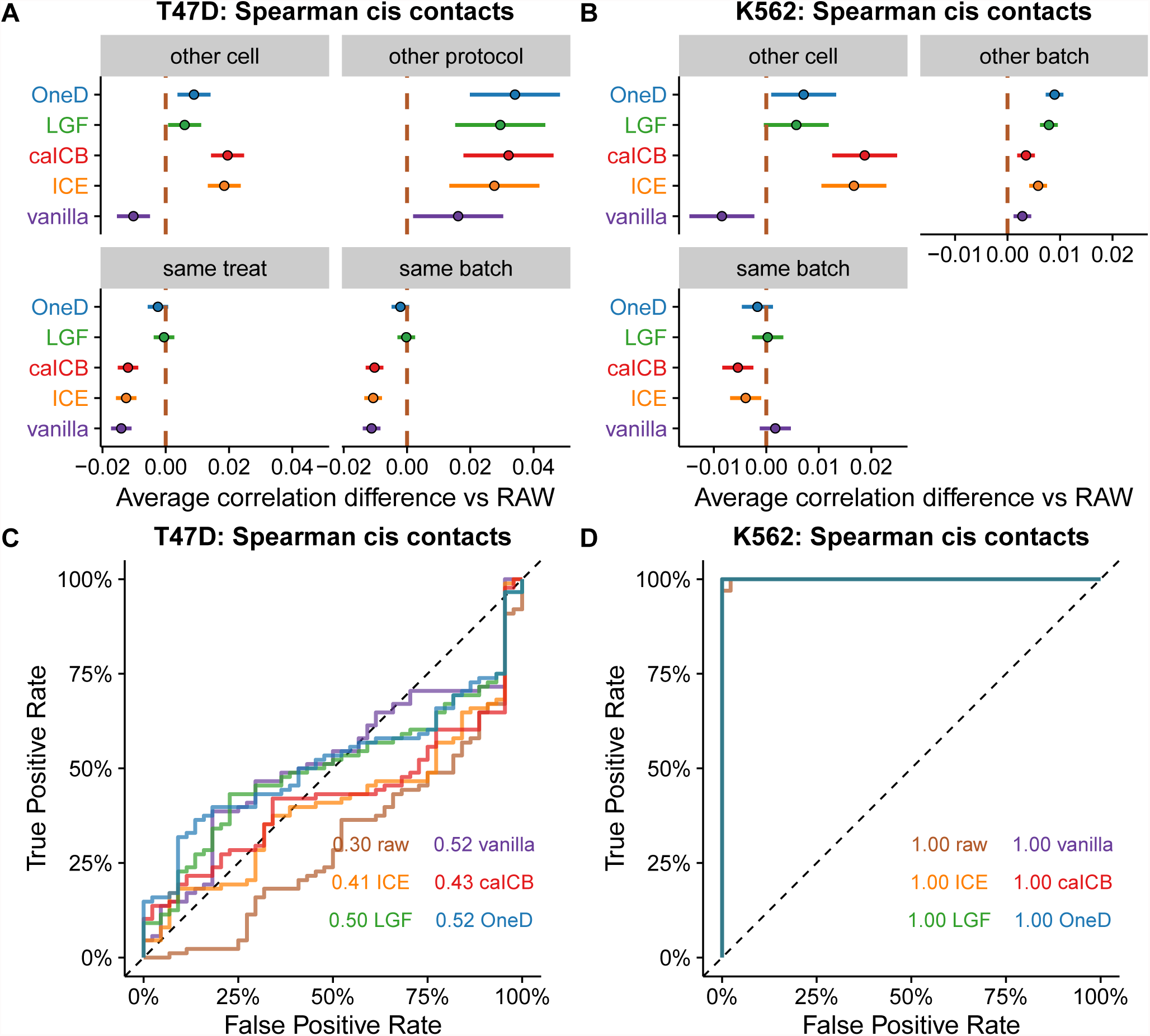
Results of the comparison between samples with aberrant karyotype. A and B. Average changes compared to raw on the T47D and K562 sets. The bars represent 95% confidence intervals centered on the mean difference of the correlation score between a given correction method and the raw data. The brown dashed line indicates the value of the average score on raw matrices (set to 0). C and D. ROC curves on the T47D and K562 sets. The areas under the curve are indicated in the bottom right corner. The color code is the same as panels A and B. The brown lines correspond to raw matrices. All results in this figure are based on Spearman correlations between the observed counts.

**Figure S5:**
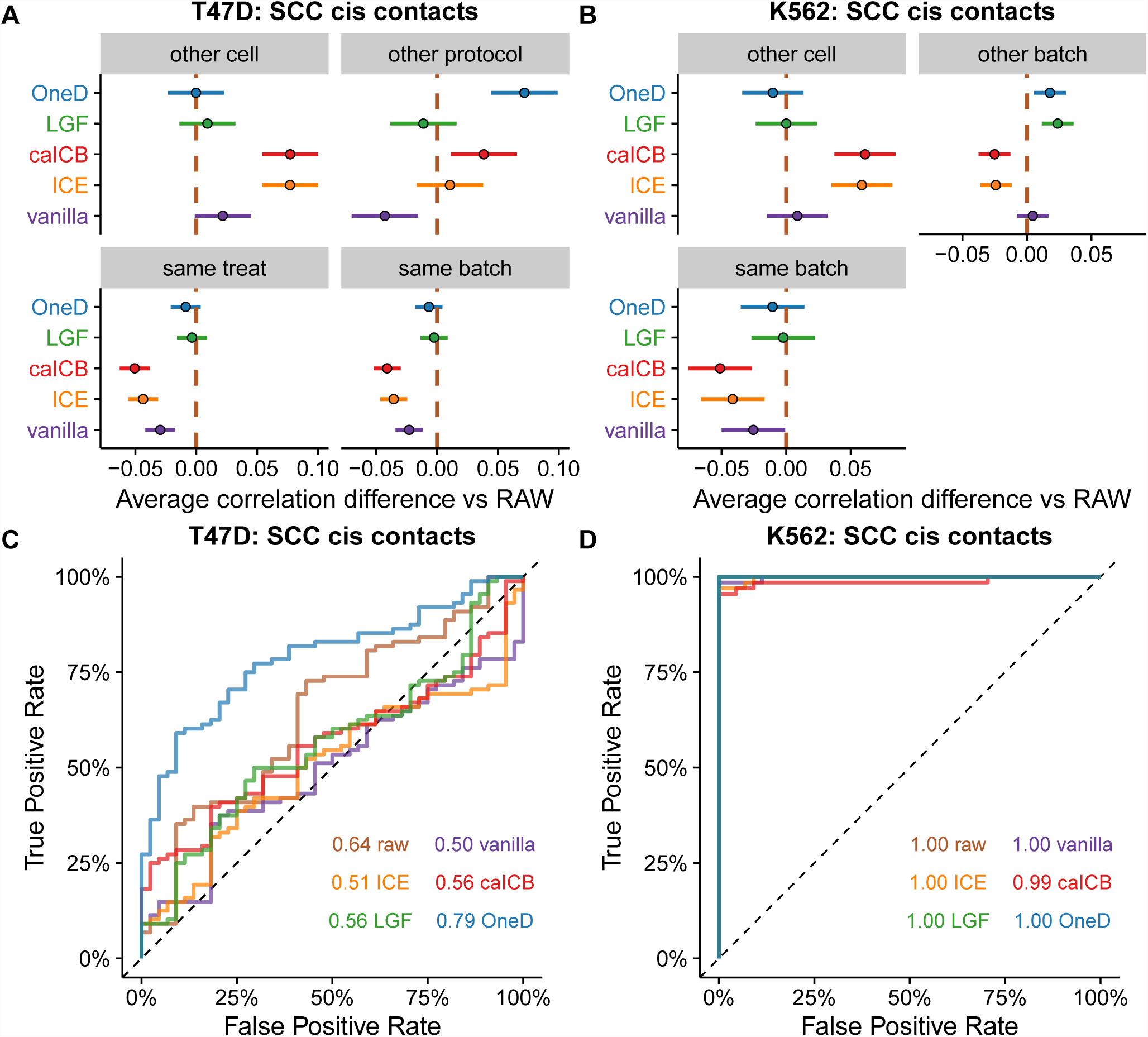
Results of the comparison between samples with aberrant karyotype. A and B. Average changes compared to raw on the T47D and K562 sets. The bars represent 95% confidence intervals centered on the mean difference of the correlation score between a given correction method and the raw data. The brown dashed line indicates the value of the average score on raw matrices (set to 0). C and D. ROC curves on the T47D and K562 sets. The areas under the curve are indicated in the bottom right corner. The color code is the same as panels A and B. The brown lines correspond to raw matrices. All results in this figure are based on stratum-adjusted correlations between the observed counts.

**Figure S6:**
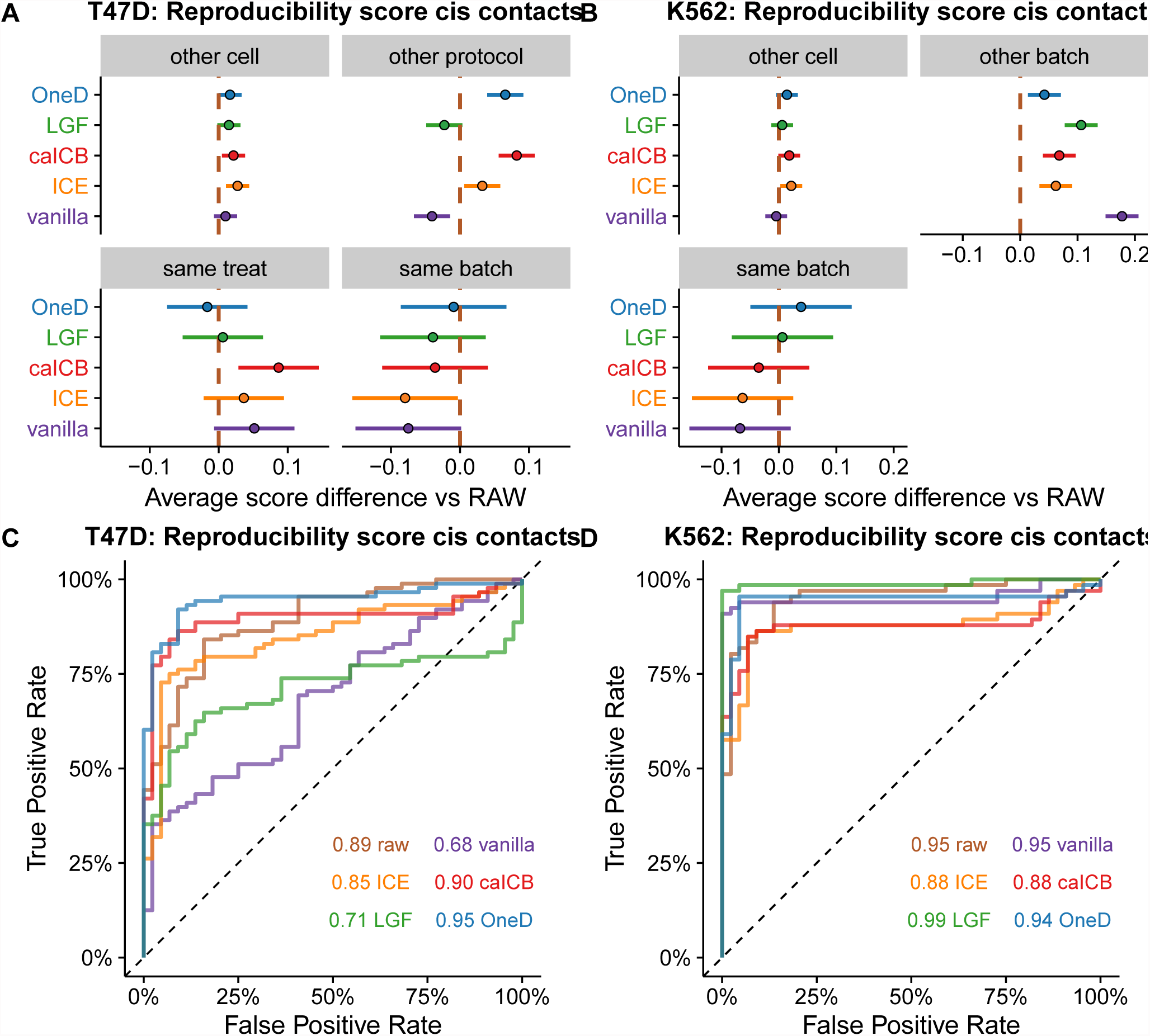
Results of the comparison between samples with aberrant karyotype. A and B. Average changes compared to raw on the T47D and K562 sets. The bars represent 95% confidence intervals centered on the mean difference of the correlation score between a given correction method and the raw data. The brown dashed line indicates the value of the average score on raw matrices (set to 0). C and D. ROC curves on the T47D and K562 sets. The areas under the curve are indicated in the bottom right corner. The color code is the same as panels A and B. The brown lines correspond to raw matrices. All results in this figure are based on the reproducibiliry score between the observed counts.

**Figure S7:**
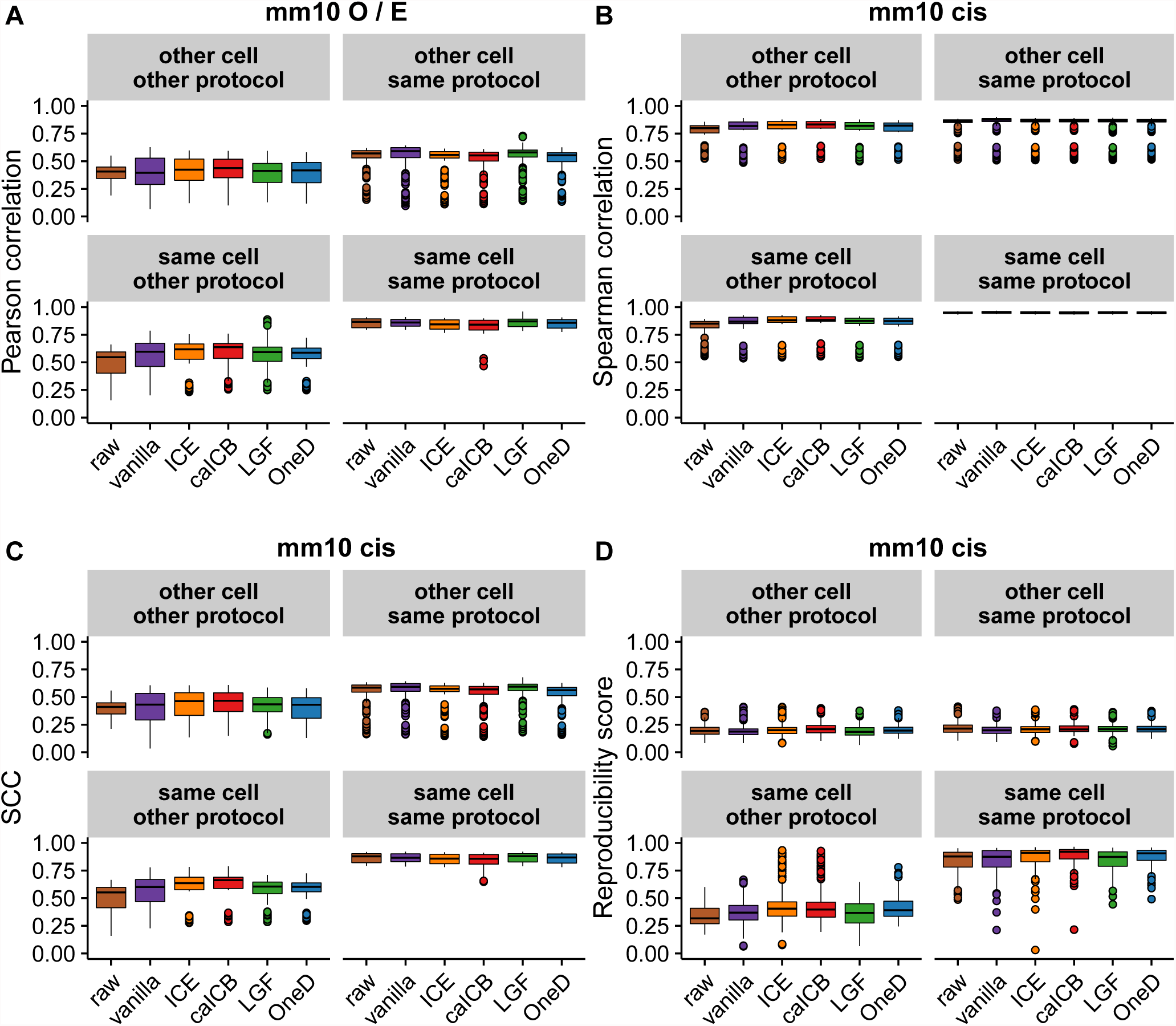
Boxplots representing the results of the pair-wise comparisons of the samples included in the mm 10 set. X axis: different correction methods. Y axis: correlation. Each panel groups paris of samples with the correspondind characteristics (in terms of cell type, protocol, batch and treatment). A. Pearson correlation between observed over expected counts. B. Spearman correlation between observed counts. C. Stratum-adjusted correlation coefficient (SCC) between observed counts. D. Reproducibility score of observed counts.

**Figure S8:**
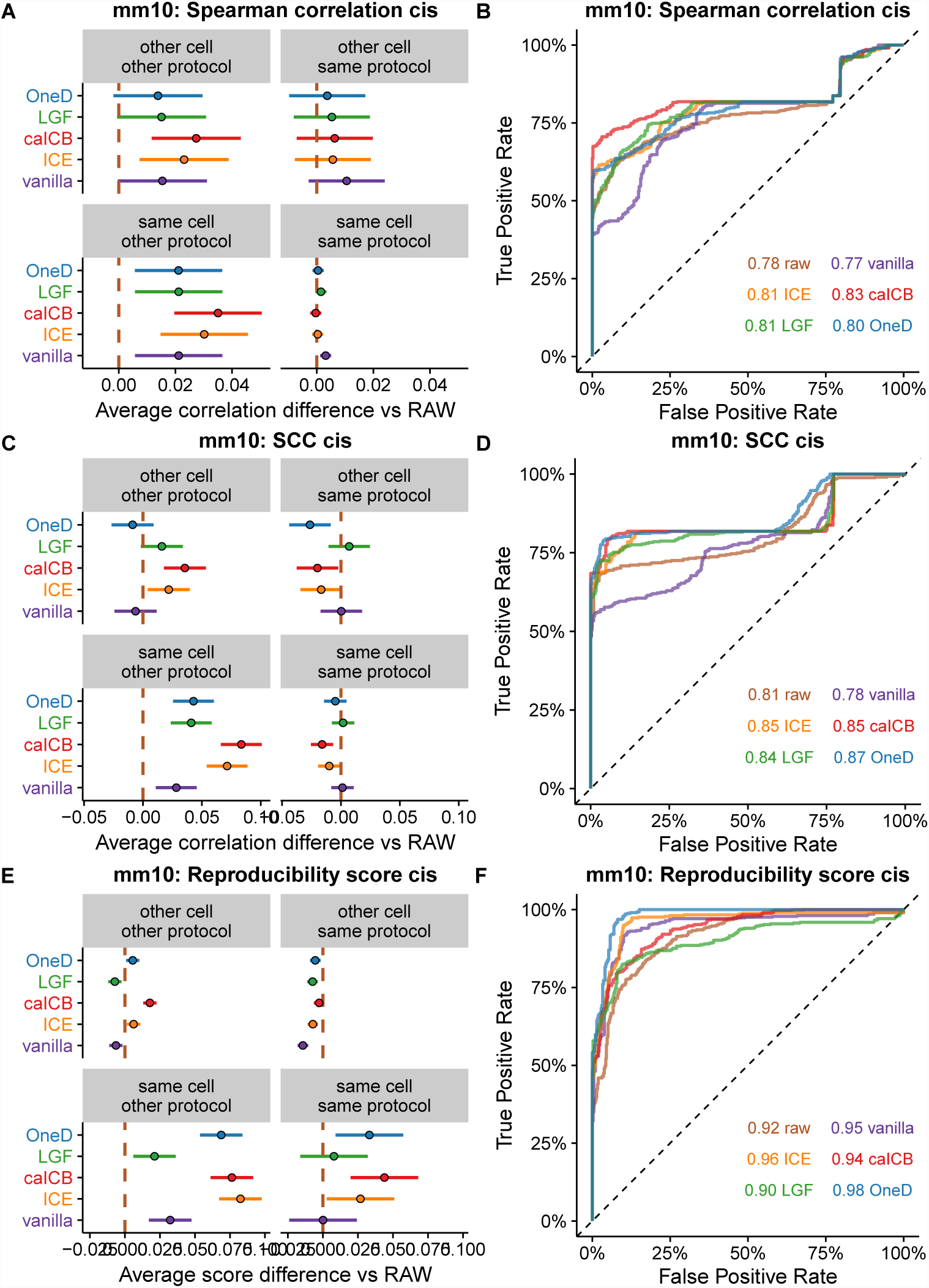
Results of the comparison between samples with normal karyotype. A., C. and E. Average changes compared to raw. The bars represent 95% confidence intervals centered on the mean difference of the correlation score between a given correction method and the raw data. The brown dashed line indicates the value of the average score on raw matrices (set to 0). B., D. and F. ROC curves. The areas under the curve are indicated in the bottom right corner. The color code is the same as panels A, C. B. The brown lines correspond to raw matrices. Results in panels A. and B. on Spearman correlations between the observed counts. Results in panels C. and D. on Stratum-adjusted correlation coefficient (SCC) between observed counts. Results in panels E. and F. on reproducibility score of observed counts.

